# BREM-SC: A Bayesian Random Effects Mixture Model for Joint Clustering Single Cell Multi-omics Data

**DOI:** 10.1101/2020.01.18.911461

**Authors:** Xinjun Wang, Zhe Sun, Yanfu Zhang, Zhongli Xu, Heng Huang, Richard H. Duerr, Kong Chen, Ying Ding, Wei Chen

## Abstract

Droplet-based single cell transcriptome sequencing (scRNA-seq) technology, largely represented by the 10X Genomics Chromium system, is able to measure the gene expression from tens of thousands of single cells simultaneously. More recently, coupled with the cutting-edge Cellular Indexing of Transcriptomes and Epitopes by Sequencing (CITE-seq), the droplet-based system has allowed for immunophenotyping of single cells based on cell surface expression of specific proteins together with simultaneous transcriptome profiling in the same cell. Despite the rapid advances in technologies, novel statistical methods and computational tools for analyzing multi-modal CITE-Seq data are lacking. In this study, we developed BREM-SC, a novel Bayesian Random Effects Mixture model that jointly clusters paired single cell transcriptomic and proteomic data. Through simulation studies and analysis of public and in-house real data sets, we successfully demonstrated the validity and advantages of this method in fully utilizing both types of data to accurately identify cell clusters. In addition, as a probabilistic model-based approach, BREM-SC is able to quantify the clustering uncertainty for each single cell. This new method will greatly facilitate researchers to jointly study transcriptome and surface proteins at the single cell level to make new biological discoveries, particularly in the area of immunology.

## INTRODUCTION

Cellular Indexing of Transcriptomes and Epitopes by Sequencing (CITE-Seq) is a recently developed revolutionary tool, which is the first technique that can measure single cell surface protein and mRNA expression level simultaneously in the same cell (1–3). Oligonucleotide-labeled antibodies are used to integrate cellular protein and transcriptome measurements. It combines highly multiplexed protein marker detection with transcriptome profiling for thousands of single cells. CITE-Seq allows for immunophenotyping of cells using existing single cell sequencing approaches (1), and it is fully compatible with droplet-based single cell RNA sequencing (scRNA-Seq) technology (e.g., 10X Genomics Chromium system (4)) and utilizes the discrete count of Antibody-Derived Tags (ADT) as the direct measurement of cell surface protein abundance. This promising and popular technology provides an unprecedent opportunity for jointly analyzing transcriptome and surface proteins at the single cell level in a cost-effective way.

In CITE-Seq experiment, the abundance of RNA and surface marker is quantified by Unique Molecular Index (UMI) and Antibody-Derived Tags (ADT) respectively, for a common set of cells at the single cell resolution. These two data sources represent different but highly related and complementary biological components. Classic cell type identification relies on cell surface protein abundance, which can be measured individually with flow cytometry. Recently, scRNA-Seq data are also used to classify cell types, based on differentially expressed genes among different cell types. In fact, both data sources have their unique characteristics and can provide complementary information. For example, the use of cell surface proteins for cell gating is advantageous in identifying common cell types but may not successfully identify some rare cell types due to its low dimensionality. On the other hand, although cell clustering based on scRNA-Seq could identify more cell types because of its higher dimensionality, it is less capable to distinguish highly similar cell types, such as CD4+ T cells and CD8+ T cells, due to a poor observed correlation between a mRNA and its translated protein expression in single cell (1,5,6).

Despite the promise of this new technology, current statistical methods for jointly analyzing data from scRNA-Seq and CITE-Seq are still unavailable or immature. A novel joint clustering approach that fully utilizes the advantages and unique features of these single cell multi-omics data will lead to a more powerful tool in identifying rare cell types or reduce false positives such as doublets. Many statistical methods have been proposed for clustering scRNA-Seq data only such as single cell interpretation via multi-kernel learning (SIMLR) (7), CellTree (duVerle, et al., 2016), Seurat (8), SC3 (9), DIMM-SC (10) and BAMM-SC (11), which are either from different clustering approach categories or recommended by recent reviewers (12,13). In contrast, to our best knowledge, there is no published method tailored for joint clustering of multi-omics data from CITE-Seq. A naïve approach is to do separate analysis on each data source, which is straightforward but suffering from various of issues such as lack of power and failing to capture the associations between transcriptome and expression of surface proteins. Multimodal data analysis, on the other hand, is supposed to achieve a more detailed characterization of cellular phenotypes than using transcriptome measurements alone.

In this study, we propose BREM-SC, a Bayesian Random Effects Mixture Model for joint clustering scRNA-Seq and CITE-Seq data. Because there is no existing method tailored for clustering single cell multi-omics data jointly, we compare the performance of BREM-SC with three popular single source clustering methods, including K-means clustering, SC3 (9) and TSCAN (14), and two commonly used multi-source clustering methods in the engineering field, including Multi-View Non-negative Matrix Factorization equipped with capped norm (MV-NMF) (15) and Pair-wised Co-regularized Multi-modal Spectral Clustering (PC-MSC) (16), in both simulation studies and real data applications. K-means is one of the most popular clustering methods and has been used in the first 10X Genomics publication (4). SC3 and TSCAN have been also proposed for clustering scRNA-Seq data, but they fall in different clustering categories. For example, SC3 is a single cell consensus clustering method, where a consensus matrix is calculated using the Cluster-based Similarity Partitioning Algorithm (CSPA). Unlike SC3, TSCAN performs model-based clustering on the transformed expression values. MV-NMF and PC-MSC are multiview extensions of two popular clustering models. MV-NMF considers the non-negative entry constraints in dimension reduction while preserving the cross-modal consistency for reduced features, after which a standard K-means is used to finalize the clustering. To make the model robust to outliers, a capped-norm objective is also utilized. Alternatively, PC-MSC introduces a co-regularization on the spectral clustering.

## MATERIAL AND METHODS

### Statistical model and estimation

The data illustration and general framework of BREM-SC modeling are shown in **Figure 1**. Although data from both sources are count data, there are several major differences between them. Firstly, the drop-out events are very common in the transcriptomic data, which are in fact much less frequent in the proteomic data. Therefore, the data matrix for RNA source is relatively sparse and people usually screen out genes with low variability in expression before performing analysis. Secondly, the overall scale of two data sets are significantly different, where proteomic data have larger values due to higher abundance of proteins in a cell. Based on the facts, we propose separate parametric model for each data source.

**Figure 1.**
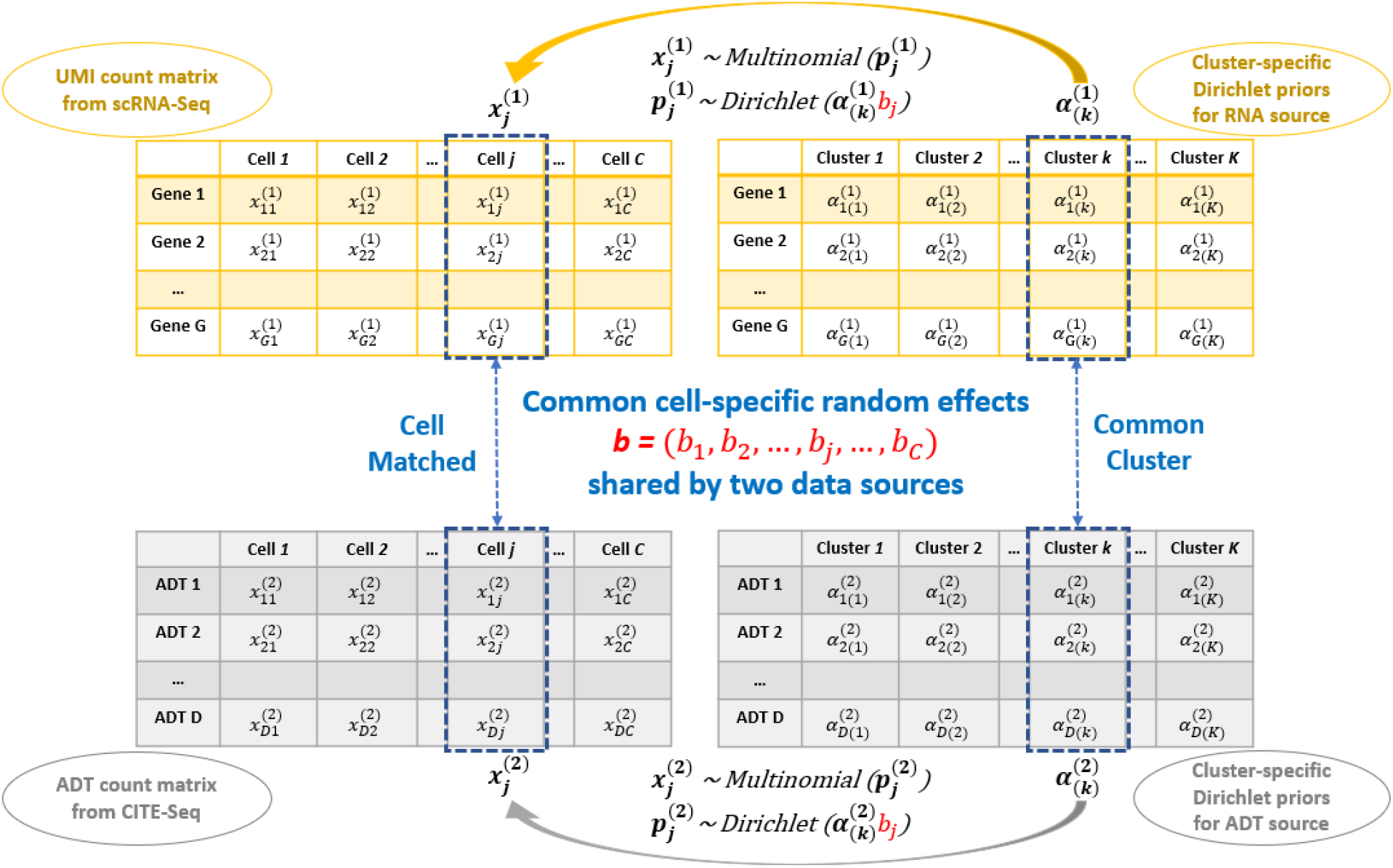
Core logic of BREM-SC method to joint clustering the RNA and ADT single cell data.

Suppose there are *C* cells generated from CITE-Seq, denote by the transcriptomic data a matrix ***X***^(**1**)^ and its ADT levels (measurement of surface protein) a matrix ***X***^(**2**)^. We use a latent variable vector ***Z*** with elements *z_j_* to represent the cell type label for cell *j*, where *j* = 1,…, *C*.

For transcriptomic data, each element 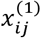 in the raw count matrix represents the number of unique UMIs for gene *i* in cell *j*, where *i* runs from 1 to the total number of genes *G*, and *j* runs from 1 to the total number of cells *C*. We then denote the number of unique UMIs in the *j* th single cell by a vector 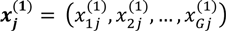. We assume that 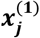 follows a multinomial distribution with parameter vector 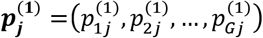. For this multinomial distribution, we further assume that the proportion 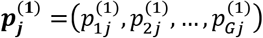 follows a Dirichlet distribution *Dir*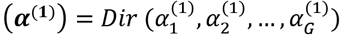, with all the elements in ***α***^(**1**)^ being strictly positive. Next, we assume that the cell population consists of *K* distinct cell types. To provide a more flexible modeling framework and allow for unsupervised clustering, we extend the aforementioned single Dirichlet prior to a mixture of *K* Dirichlet distributions, indexed by *k* = 1, …,*K* and each with parameter 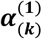. For instance, if cell *j* belongs to the *k* th cell type, its gene expression profile 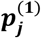 follows a cell-type-specific prior distribution 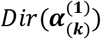. The Dirichlet multinomial density for cell *j*, as illustrated in DIMM-SC (10), can be obtained by multiplying the Dirichlet mixture prior by the multinomial density and then integrating out 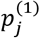, which is derived as 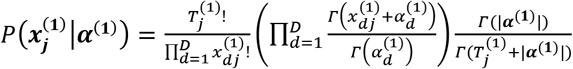, where 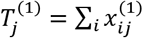 is the total number of unique UMIs for *j* th cell.

Similarly, we also use the Dirichlet multinomial distribution to model surface protein (ADT) data. Suppose there are in total *D* number of ADT markers, then similarly, the density of Dirichlet multinomial is derived as 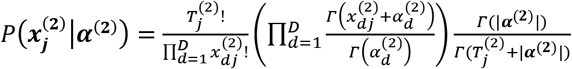, where *d* = 1,…,*D* indicates the index for protein marker and 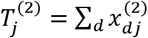 is the total ADT counts for *j* th cell.

Without considering the correlation between two datasets, the joint distribution for all cells, assuming full independency between two sources, can then be derived as

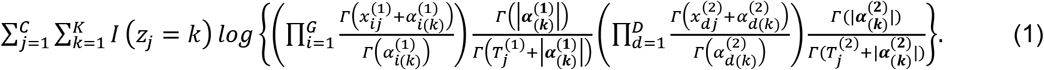

To further model the correlation between different data sources, we introduce cell-specific random effects into our framework. Since we assume that cells belonging to same cell type share common Dirichlet parameters, we model cell heterogeneity by directly multiplying cluster-specific Dirichlet parameters with the random effects, i.e. 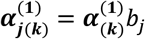 and 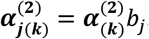, where 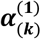 and 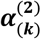 are the Dirichlet parameters of cell type *k* for RNA and protein data, respectively, and *b_j_* is the random effects for the *j* th cell. Based on the fact that Dirichlet parameters are strictly positive numbers, we assume that this cell-specific random effects *b_j_* ~ *LogNormal* 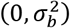, i.e., 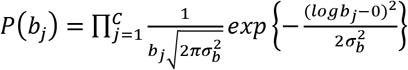. Finally, the complete log likelihood can be derived as

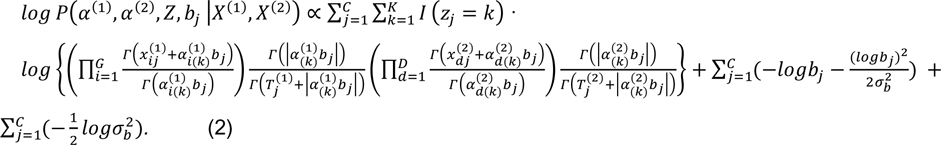

We use Gibbs sampler to iteratively update 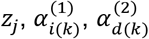, and *b_j_* in Equation 2 (Details can be found in **Supplementary Methods**).

### Selection of the number of clusters and initial values

To implement BREM-SC, it is critical to select the total number of clusters and the initial values for MCMC. Specifically, the number of clusters *K* can be defined either with prior knowledge or standard model checking criterion such as Akaike’s Information Criteria (AIC) or Bayesian Information Criteria (BIC). Meanwhile, there are many methods to determine the initial values of *α*_1_,*α*_2_, …,*α_G_*. For BREM-SC we applied K-means clustering to get a preliminary clustering result for each data source separately, followed by using Ronning’s method (17) to estimate initial values of ***α***, which is similar to the estimation procedure implemented in DIMM-SC (10).

### Data generation algorithm for simulation studies under BREM-SC model

Based on different parameter settings in our Dirichlet multinomial models, we simulated RNA expression and ADT measurements for each single cell. In the simulation set-up, the two count matrices were sampled from the proposed Dirichlet mixture models. Specifically, for a fixed number of cell clusters *K*, we first pre-defined the values of 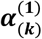 and 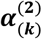 for the *k* th cell cluster. The random effects *b_j_* are generated from a log-normal distribution with pre-specified value 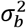. We then computed the transcriptomic profile 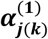 for each single cell by multiplying 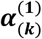 by *b_j_*. Similarly, for cellular protein expression profile, we multiplied 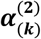 by *b_j_* to compute 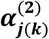. For the next step, we sampled the cluster proportion 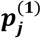 (or 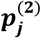) from a Dirichlet distribution *Dir* 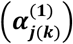 (or *Dir* 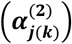). Lastly, we sampled the UMI count vector 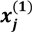 for *j* th cell from multinomial distribution *Multinomial*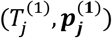 and sampled the ADT count vector 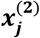 from *Multinomial*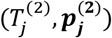, where 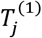 and 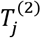 are each simulated from truncated normal distribution with parameters estimated from real data.

### Data generation algorithm for simulation studies under model misspecification

We used R package *Splatter* (18) to simulate data to assess robustness of BREM-SC under model misspecification. In *Splatter,* the final data matrix is a synthetic dataset consisting of counts from a Gamma-Poisson (or negative-binomial) distribution. Since there is no existing method for generating single cell surface protein expression levels from CITE-Seq, we also used *Splatter* to generate ADT count for the proteomic data. To make our simulated gene expression data a good approximation to the real data, our model parameters (in *Splatter)* were estimated from the real data downloaded from 10X Genomics website (https://support.10xgenomics.com/single-cell-gene-expression/datasets/1.1.0/b_cells). Specifically, for ADT data, we modified the *Splatter* parameters such as dropout rate, library size, expression outlier, and dispersion across features to make the simulated data more similar to real observed ADT data regarding the scale. We assumed all cell types are shared between gene expression and ADT data, and further specified differential expression parameters to generate scenarios with different magnitude of cell type differences.

### Setup of competing methods used in this paper

In general, all clustering methods to which BREM-SC compared were performed under their default parameters. Single-source clustering methods including K-means, SC3 and TSCAN were applied to the pooled data from RNA expression and ADT with centered log-ratio (clr)-transformation (8) for normalization while ignoring source specificity.

### Metric for clustering performance

We used adjusted rand index (ARIs), a commonly used metric to indicate similarity between two clustering results (19). ARI ranges from −1 to 1. An ARI of value 1 indicates the two clustering results are perfectly concordant, while 0 indicates a result by random clustering. On the other hand, an ARI of negative value corresponds to a situation in which the clustering result is even worse than randomly assigning clusters.

### Public human peripheral blood mononuclear cells (PBMC) CITE-Seq dataset

To assess the performance of BREM-SC, we used a published human PBMC CITE-Seq dataset downloaded from 10X Genomics website (https://support.10xgenomics.com/single-cell-gene-expression/datasets/3.0.0/pbmc_10k_protein_v3?src=search&lss=google&cnm=sem-goog-2020-website-page-ra_g-p_brand-amr-retarget&cid=7011P000000oWys). A total of 7,865 cells from a healthy donor were stained with 14 TotalSeq-B antibodies, including CD3, CD4, CD8a, CD14, CD15, CD16, CD19, CD25, CD45RA, CD45RO, CD56, CD127, PD-1 and TIGIT. Matched scRNA-Seq data are available.

### In-house human PBMC CITE-Seq dataset

To further evaluate the validity of our method. We generated an in-house CITE-seq dataset of human PBMC from a healthy. 1,372 cells were stained with Totalseq-A and are prepared using the 10x Genomics platform with Gel Bead Kit V2. The prepared assay is subsequently sequenced on an Illumina Hiseq with a depth of 50K reads per cell. Cells in this dataset are measured for their surface marker abundance through CITE-seq (1). Ten surface markers are measured for every cell: CD3, CD4, CD8a, CD11c, CD14, CD16, CD19, CD56, CD127 and CD154.

## RESULTS

To assess the performance of BREM-SC, we performed comprehensive simulation studies to compare BREM-SC with five existing clustering methods, including K-means, SC3, TSCAN, MV-NMF and PC-MSC, all of which are hard clustering approaches and thus will assign each cell to an exclusive cluster. We simulated scRNA-Seq and CITE-Seq data under both our proposed model and model misspecification (using R package Splatter (18)) to assess the validity and robustness of BREM-SC. For each simulation scenario, we simulated 100 datasets to assess the variability of clustering results for each method. We also applied BREM-SC on two human peripheral blood mononuclear cells (PBMC) CITE-Seq datasets to assess the usefulness of our method in real application. In general, we used adjusted rand index (ARI) as the metric to assess the clustering performance.

### BREM-SC outperforms other methods in simulation studies under our proposed model

We designed different simulation scenarios to assess the performance of BREM-SC by varying number of cells in each cluster, number of clusters, magnitude of cell type differences (i.e., the magnitude of difference among different clusters), and among-cell variabilities (indicated by the magnitude of random effects).

The results of simulation studies under our proposed model are shown in **Figure 2**. In general, the performance of all clustering approaches decreases as the among-cell variability, indicated by 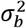, increases, number of clusters increases, and number of cells decreases. As shown in **Figure 2A**, BREM-SC outperformed the other five competing methods by achieving the highest average ARI among 100 simulations when the level of among-cell variabilities is medium or large. However, when 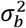 is small (i.e., *b_j_*’s are all close to 1) which indicates an extreme case in our random effects framework, BREM-SC, although achieves highest median ARI, demonstrated larger variability compared to other methods.

**Figure 2.**
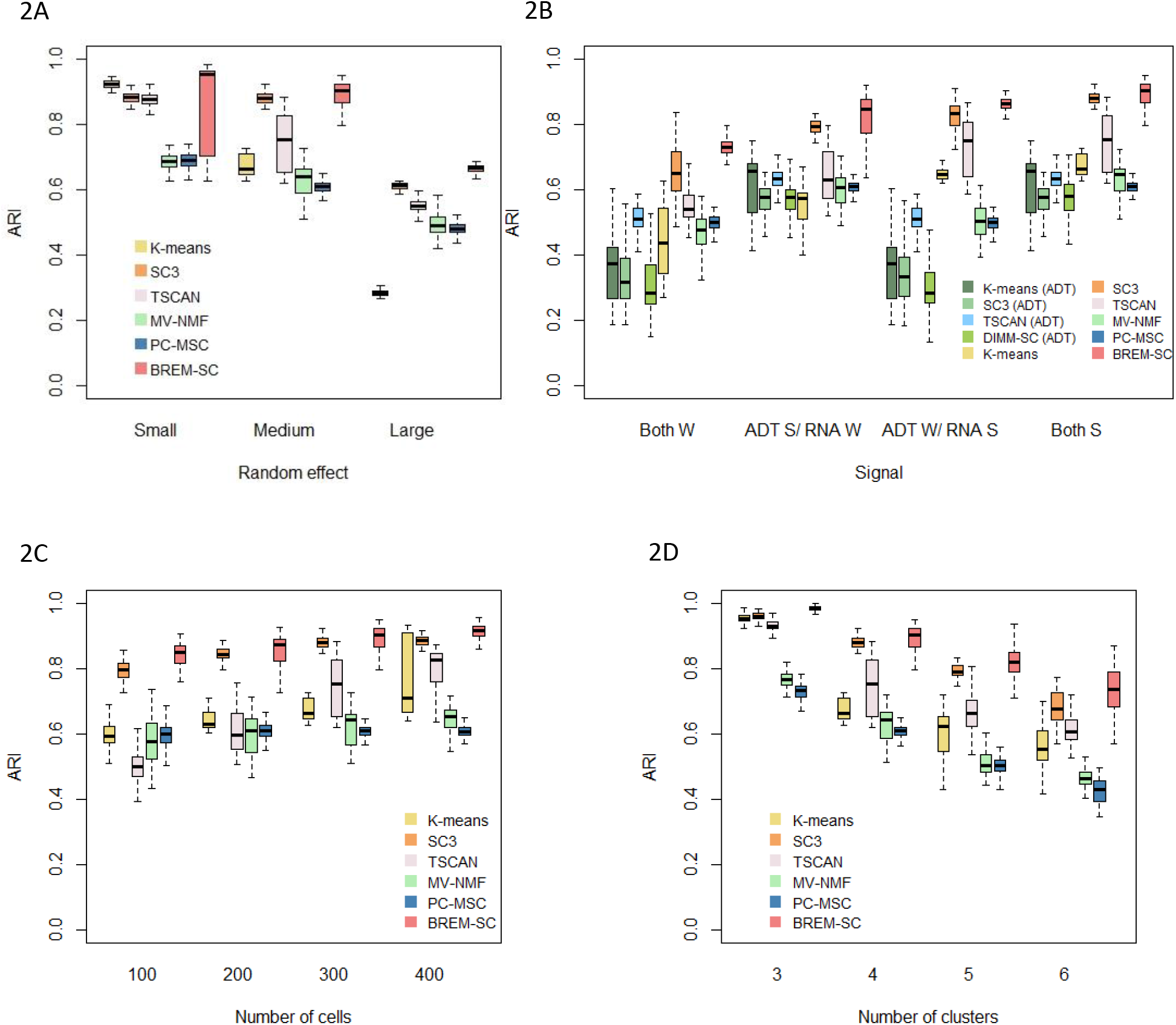
Boxplots of ARIs for six clustering methods across 100 simulations, investigating how various among-cell variabilities (**2A**), magnitude of cell type differences (**2B**, W refers to weak signals between clusters; S refers to strong signals), number of cells in each cluster (**2C**) and number of clusters (**2D**) affect the clustering results.

**Figure 2B–2D** list the boxplots of ARI by varying magnitude of cell type differences, number of cells in each cluster, and number of clusters, respectively. In **Figure 2B**, we considered four scenarios in terms of signal strength from two data sources. To illustrate the advantage of joint clustering, we also applied K-means, SC3, TSCAN and DIMM-SC on ADT data alone. When the clustering signal is strong (i.e., difference among cell clusters is large) in both RNA expression and ADT data (referred to *Both S* column), both BREM-SC and SC3 performed extremely well while other methods show fair clustering results. However, when cell clusters are similar in either proteomics (referred to *ADT W/RNA S* column) or transcriptomics data (referred to *ADT S/RNA W* column), K-means and TSCAN produced less accurate clustering results, while BREM-SC and SC3 still performed well. As expected, strong clustering signal leads to higher clustering accuracy and lower clustering variability. If both RNA expression and ADT data are alike across different cell types (referred to *Both W* column), ARIs of all methods decreased but BREM-SC still performed the best. This simulation scenario clearly demonstrated that as long as either of the data source contains strong clustering signals, our BREM-SC takes full advantage of that and achieves highly satisfactory performance. In **Figure 2C**, we observed that more cells can provide more accurate and robust clustering results, which is as expected. In **Figure 2D**, we found that larger number of clusters is associated with worse clustering performance. Again, in both scenarios BREM-SC outperformed the other five methods across all various settings.

Consistently across all four scenarios shown in **Figure 2**, when data are generated from our proposed true model, BREM-SC outperformed K-means clustering, SC3, TSCAN, MV-NMF and PC-MSC, suggesting its advantages in fully utilizing both types of data simultaneously.

### BREM-SC outperforms other methods in simulation studies under model misspecification

To evaluate the robustness of BREM-SC when the data generation model is mis-specified, we simulated additional datasets using R package *Splatter* (20), a commonly used tool to simulate scRNA-Seq data whose underlying models are completely different from ours.

**Figure 3** shows the performance of BREM-SC and other competing methods under model misspecification. In **Figure 3A**, we simulated different signal strength for each data source using *Splatter.* Methods such as K-means and MV-NMF produced poor results across all scenarios even when signal strengths are large in both data sources. On the other hand, BREM-SC outperformed all other methods and showed good clustering performance even when signal strengths are relatively small in both data sources. In **Figure 3B**, we assessed the clustering performance across different probabilities that a gene is differentially expressed (DE). In general, higher DE probability indicates easier cell clustering. Again, we found that BREM-SC outperformed all other competing methods in terms of clustering accuracy. Therefore, we demonstrated the strong robustness of BREM-SC under model misspecification.

**Figure 3.**
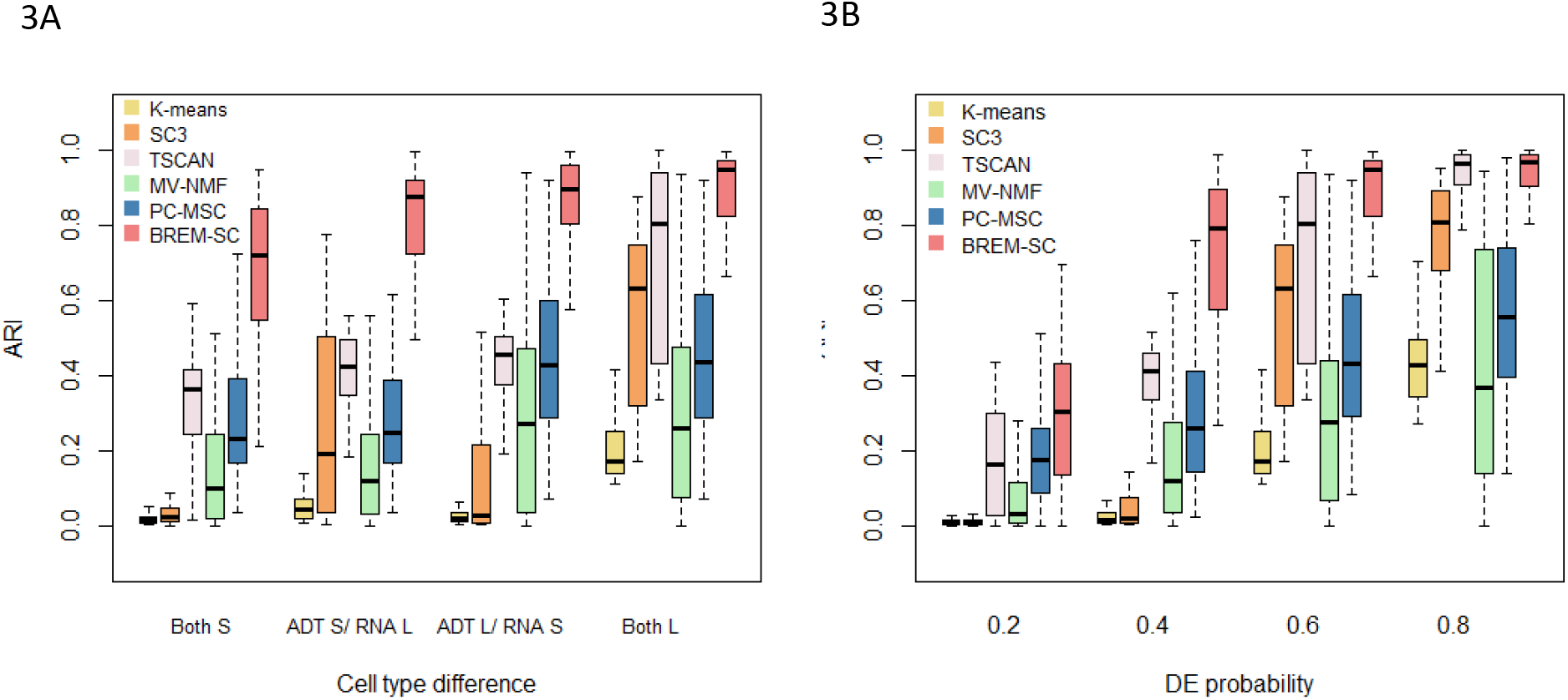
Boxplots of ARIs for six clustering methods across 100 simulations using Splatter. The performance under various magnitude of cell type differences were investigated. In Figure 3A, mean parameters of three cell types were set as (0.15, 0.151, 0.152) and (0.15, 0.2, 0.25) to represent two levels (S standards small and L standards for large) of cell type difference. In Figure 3B, different probabilities that a gene would be selected to be differentially expressed (DE) were set.

### Analysis of a public human PBMC CITE-Seq dataset

To evaluate the clustering performance of BREM-SC on real data, we first used a published human PBMC CITE-Seq dataset downloaded from 10X Genomics website. A total of 7,865 cells and 14 surface protein markers are included in this dataset in addition to matched scRNA-Seq data. As a standard approach of analyzing human PBMC dataset, we extracted the top 100 highly variable genes based on the rank of their standard deviations in the scRNA-Seq data. We identified seven cell types based on the biological knowledge of both protein and gene markers as the approximate truth. Examples of such cell type identification procedure are shown in **Supplementary Figure S1**. Taken together, more than 80% of single cells can be assigned to a specific cell type. Cells with uncertain cell types (not identified in the ground truth) were removed from computing ARIs.

We applied seven clustering methods (K-means clustering, TSCAN, SC3, DIMM-SC, MV-NMF, PC-MSC and BREM-SC) on this dataset and performed both single source clustering (using ADT or RNA data only) and multi-source joint clustering, respectively. We ran each method ten times to evaluate the stability of clustering performance. Note that since TSCAN is a deterministic clustering method and generates identical results, stability of performance cannot be assessed from analyzing a single dataset for this method. As shown in **Table 1**, BREM-SC outperformed other methods for multi-source joint clustering in terms of ARI. In addition, because the ground truth is built based on both protein and mRNA information, BREM-SC, as a joint clustering method utilizing both information appropriately, produced better results compared to all single source clustering analyses in this application. Further, we used Uniform Manifold Approximation and Projection (UMAP) plot (21) to visualize the clustering results from BREM-SC. UMAP is a recently developed non-linear dimension reduction tool for visualization, but quickly gains popularity among single cell researches because of its outstanding performance. **Figure 4** shows the UMAP plots with each cell colored by their ground truth label (**Figure 4A**) and cluster labels inferred by BREM-SC (**Figure 4B**), respectively. In general, the two plots are highly similar regarding to the distribution of different clusters (ARI = 0.868), indicating the outstanding performance of BREM-SC.

**Table 1.** Performance (ARI) of different clustering methods from ten times analyses of the public human PBMC real dataset

|  | Mean (SD) | Median (Range) |
| --- | --- | --- |
| Single source clustering - RNA only |  |  |
| K-means | 0.451 (0.066) | 0.487 (0.335, 0.536) |
| TSCAN | 0.458 (N/A) | 0.458 (N/A) |
| SC3 | 0.505 (0.070) | 0.528 (0.366, 0.571) |
| DIMM-SC | 0.529 (0.025) | 0.526 (0.503, 0.555) |
| Single source clustering - protein only |  |  |
| K-means | 0.653 (0.093) | 0.648 (0.436, 0.760) |
| TSCAN | 0.389 (N/A) | 0.389 (N/A) |
| SC3 | 0.649 (0.045) | 0.668 (0.568, 0.692) |
| DIMM-SC | 0.673 (0.048) | 0.679 (0.618, 0.737) |
| Multi-source joint clustering |  |  |
| K-means | 0.645 (0.158) | 0.689 (0.383, 0.889) |
| TSCAN | 0.472 (N/A) | 0.472 (N/A) |
| SC3 | 0.543 (0.130) | 0.534 (0.395, 0.853) |
| MV-NMF | 0.685 (0.1) | 0.671 (0.546, 0.856) |
| PC-MSC | 0.564 (0.037) | 0.547 (0.523, 0.61) |
| BREM-SC | 0.737 (0.125) | 0.713 (0.556, 0.868) |

### “Soft-clustering” property of BREM-SC

We further used this public human PBMC CITE-Seq dataset to illustrate the property of “soft-clustering”, that BREM-SC can provide the posterior probability that a given cell belongs to a specific cluster in addition to cell labels. We highlighted the “vague” cells identified by BREM-SC on UMAP plots. Here “vague” cells refer to those cells with largest posterior probability smaller than a pre-specified threshold. By setting up different probability threshold, the number of “vague” cells can be controlled. In **Figure 5** we showed the distribution of “vague” cells on UMAP plot for the public PBMC dataset varying by different threshold. By comparing with **Figure 4A**, most of such “vague” cells coincide with either the unknown cells that we fail to identify based on current biological knowledge or the cells on the boundary between two cell types that are closely attached to each other on UMAP plot.

**Figure 4.**
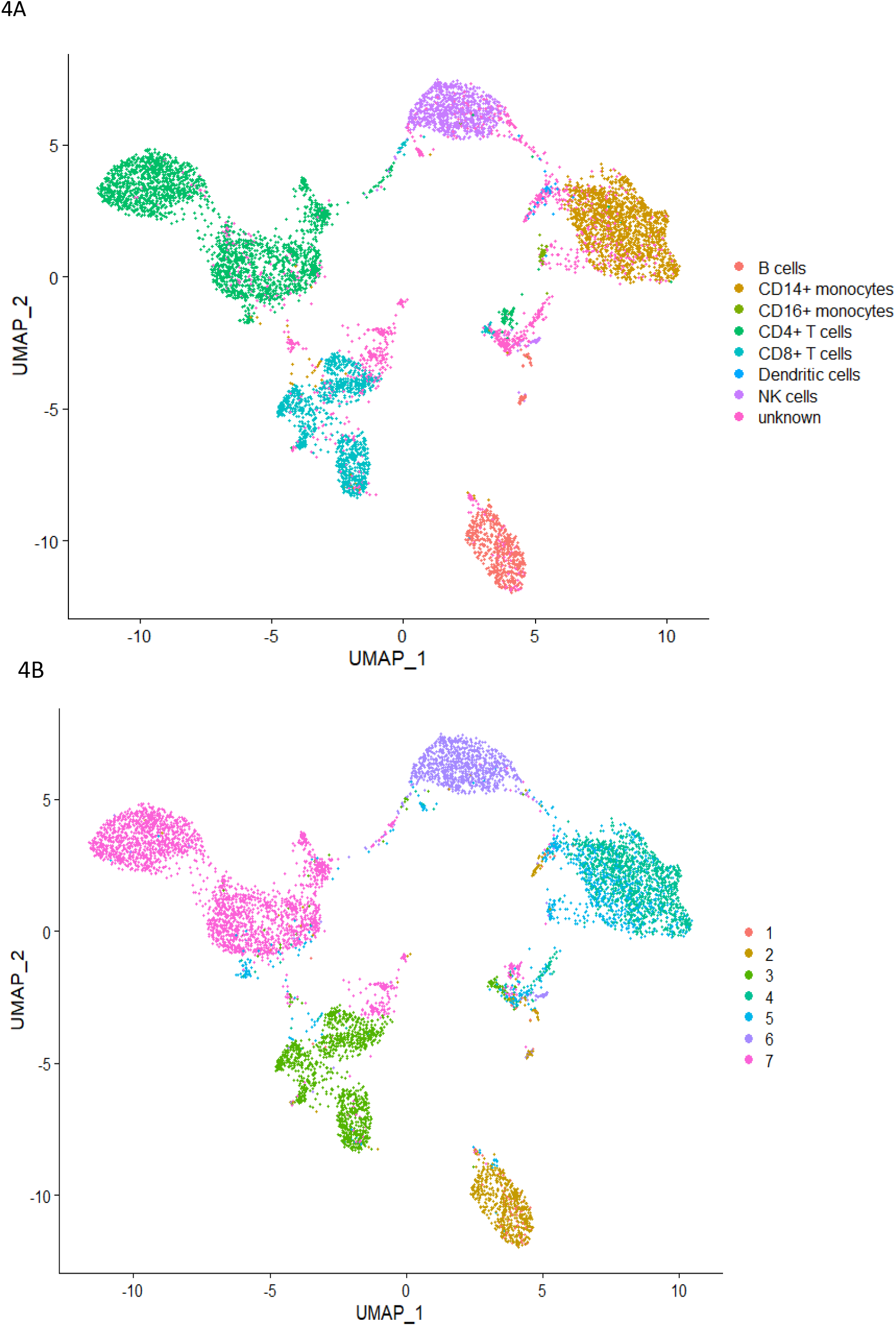
The performance of BREM-SC with the public human PBMC CITE-Seq dataset (from 10X Genomics). The UMAP projection of cells are colored by the ground truth (**4A**) and BREM-SC clustering results (**4B**).

**Figure 5.**
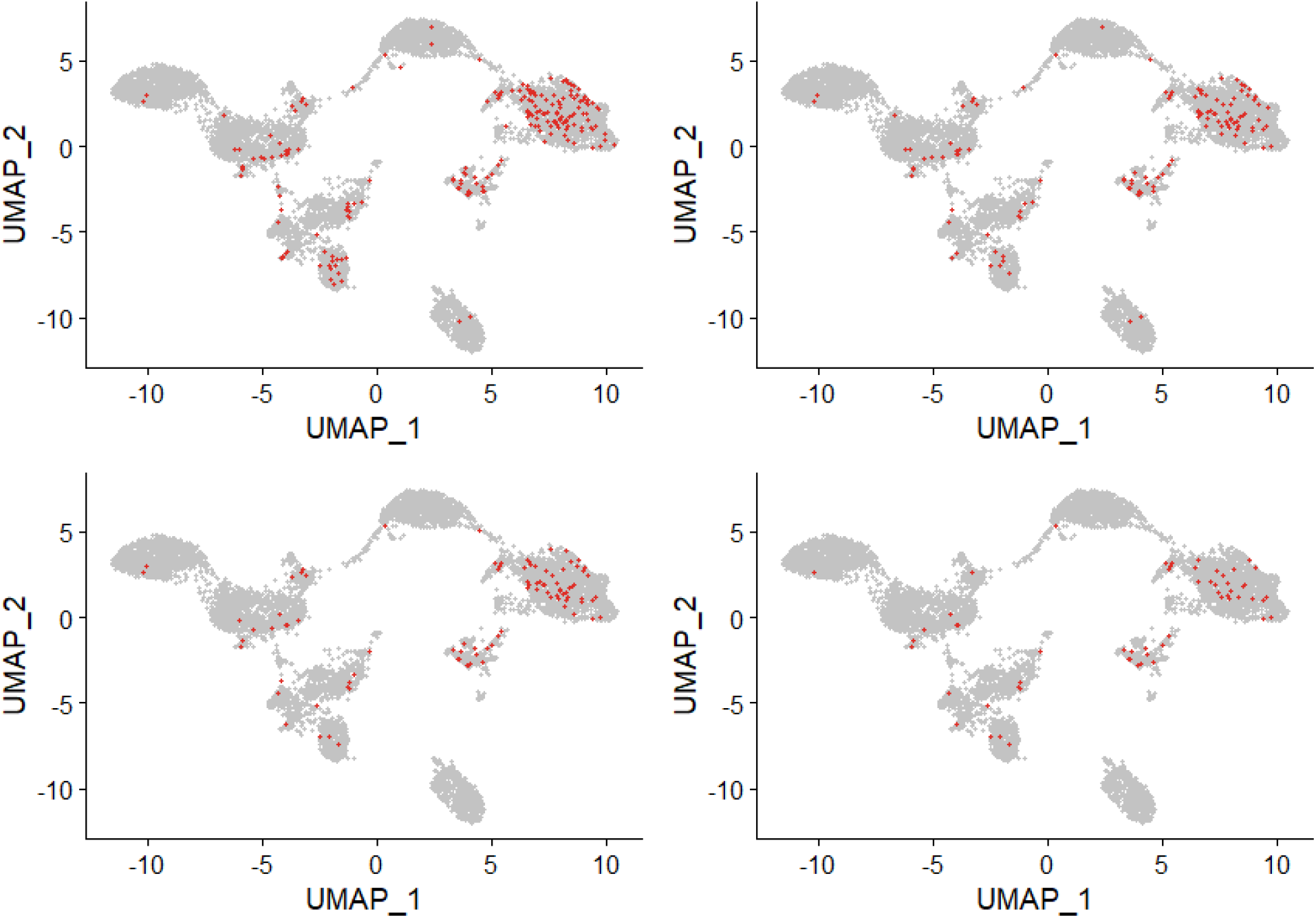
Illustration of “soft clustering” property by highlighting vague cells with the largest posterior probability < 0.99 (top left); < 0.95 (top right); < 0.9 (bottom left); and < 0.8 (bottom right) from BREM-SC.

### Analysis of an in-house human PBMC CITE-Seq dataset

Because there are limited public datasets of CITE-seq, to further evaluate our method, we performed a CITE-seq experiment on human PBMCs from a healthy donor with informed consent and University IRB approval. 10X Genomics Chromium system was used to generate CITE-Seq data with 10 cell surface markers designed in the experiment, which yielded a total of 1,372 cells. Similarly, we extracted the top 100 highly variable genes based on their standard deviations in the RNA data. For this dataset, we identified six subtypes of PBMCs based solely on the biological knowledge of cell-type-specific protein markers, and more than 85% of single cells were assigned to a specific cell type using these markers. Examples of such cell type identification procedure are shown in **Supplementary Figure S2**. We then used these cell labels as the approximated ground truth to benchmark the clustering performance for different clustering methods. Cells with uncertain cell types were removed when calculating ARIs.

Similar to the analysis of public human PBMC dataset, we applied seven clustering methods on this inhouse PBMC dataset and repeated each method ten times to evaluate the stability of its performance. As shown in **Figure 6**, BREM-SC performed very well in the human PBMC samples, since the UMAP plot with each cell colored by their cell-type label based on protein markers (**Figure 6A**) is highly similar to the plot generated from the clustering result of BREM-SC (**Figure 6B**). Clustering results for all methods are summarized in **Table 2**. Again, BREM-SC outperformed all other competing methods for multi-source joint clustering based on the average ARI. However, it is observed that in general single source clustering with ADT count data performed better than multi-source joint clustering under this circumstance. The reason for this result is that the approximated ground truth is built solely based on protein markers, and thus will favor clustering analyses only using the protein data. Methods such as K-means and TSCAN performed much worse compared to their corresponding single source analysis results, indicating that they failed to properly incorporate RNA information and thus the joint clustering results were highly compromised by the inclusion of RNA “noises”. On the other hand, BREM-SC, although performed slightly worse than DIMM-SC with ADT only clustering, still gives satisfactory results (mean ARI = 0.857), indicating that BREM-SC can handle mRNA-protein association very well in real data and is robust to “noises” introduced from one data source.

**Figure 6.**
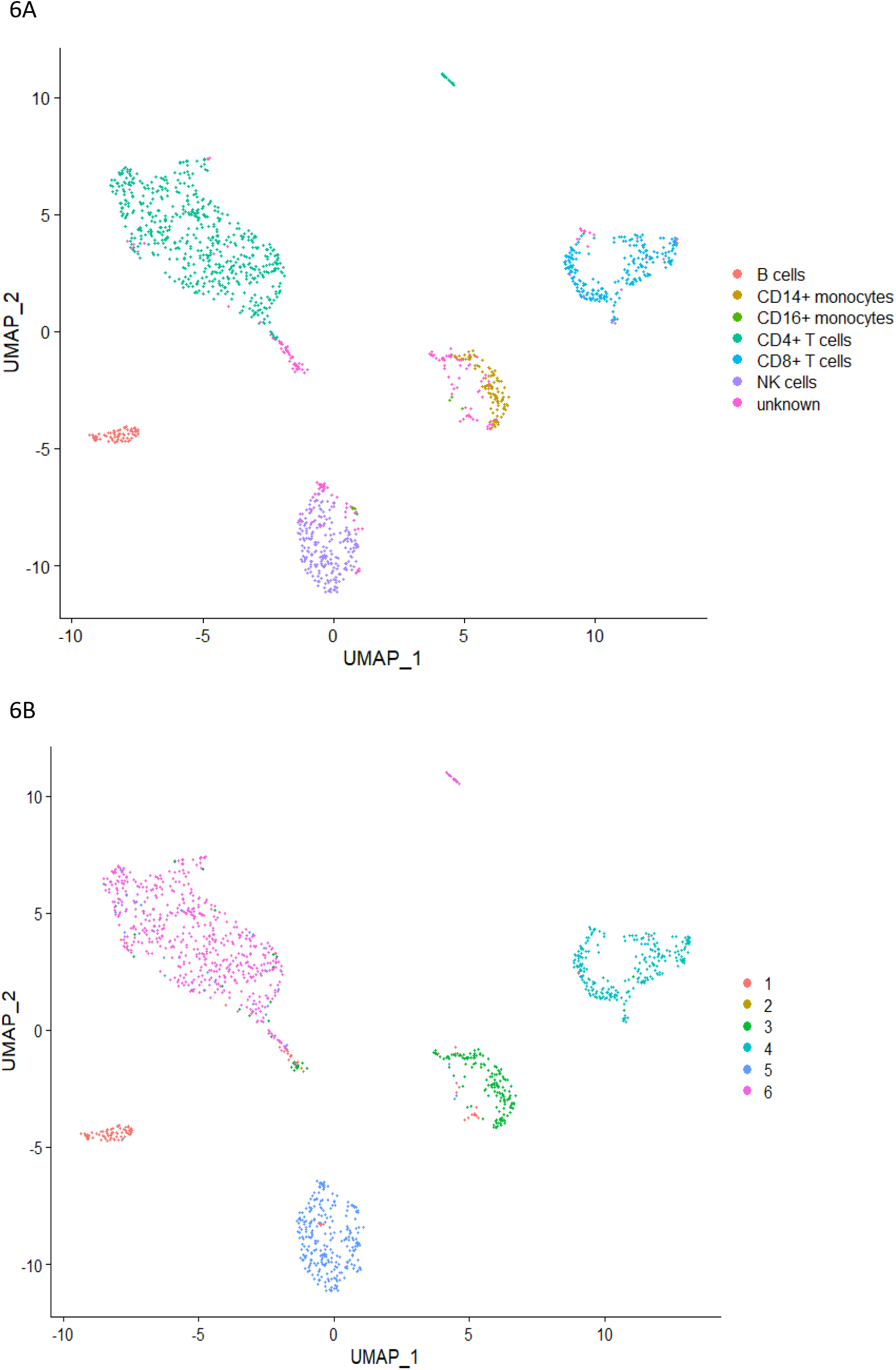
The performance of BREM-SC for in-house human PBMC CITE-Seq dataset. The UMAP projection of cells are colored by the ground truth (**6A**) and BREM-SC clustering results (**6B**).

**Table 2.** Performance (ARI) of different clustering methods from ten times analyses of the in-house human PBMC real dataset

|  | Mean (SD) | Median (Range) |
| --- | --- | --- |
| Single source clustering - RNA only |  |  |
| K-means | 0.304 (0.041) | 0.295 (0.275, 0.415) |
| TSCAN | 0.343 (N/A) | 0.343 (N/A) |
| SC3 | 0.381 (0.037) | 0.383 (0.321, 0.438) |
| DIMM-SC | 0.401 (0.011) | 0.407 (0.380, 0.409) |
| Single source clustering - protein only |  |  |
| K-means | 0.930 (0.120) | 0.986 (0.702, 0.989) |
| TSCAN | 0.877 (N/A) | 0.877 (N/A) |
| SC3 | 0.921 (0.039) | 0.931 (0.816, 0.951) |
| DIMM-SC | 0.959 (0.037) | 0.985 (0.916, 0.992) |
| Multi-source joint clustering |  |  |
| K-means | 0.596 (0.142) | 0.613 (0.434, 0.803) |
| TSCAN | 0.443 (N/A) | 0.443 (N/A) |
| SC3 | 0.777 (0.047) | 0.788 (0.679, 0.848) |
| MV-NMF | 0.735 (0.118) | 0.701 (0.554, 0.942) |
| PC-MSC | 0.612 (0.086) | 0.646 (0.502, 0.69) |
| BREM-SC | 0.857 (0.048) | 0.874 (0.749, 0.895) |

### Realistic simulation studies with in-house human PBMC CITE-Seq dataset

In practice, it is common that researchers may have limited biological information of a specific cell type (especially rare cell type). As a result, they may fail to include some protein markers that are specifically differentially expressed in that rare cell type when they design their CITE-Seq experiments. In this case, researchers are not able to identify all the cell types with surface protein data only, and joint clustering methods will be preferred if they can appropriately incorporate RNA data to compensate the “loss” of surface markers. To mimic the situation where the pre-designed protein markers cannot capture the characteristics of all cell types, we removed three protein markers (CD8A, CD16, CD127) in this human PBMC dataset and re-analyzed the remaining data. The results of this realistic simulation are summarized in **Table 3**. In general, the results of single source clustering analyses are much worse when some important protein markers are missing. BREM-SC, on the other hand, achieves the highest mean ARI compared with all other clustering methods (both single source and multi-source). In addition, the performance of BREM-SC is only slightly influenced with missing a few protein markers, which demonstrates the robustness of BREM-SC and its capability of properly handling the mRNA-protein association.

**Table 3.**
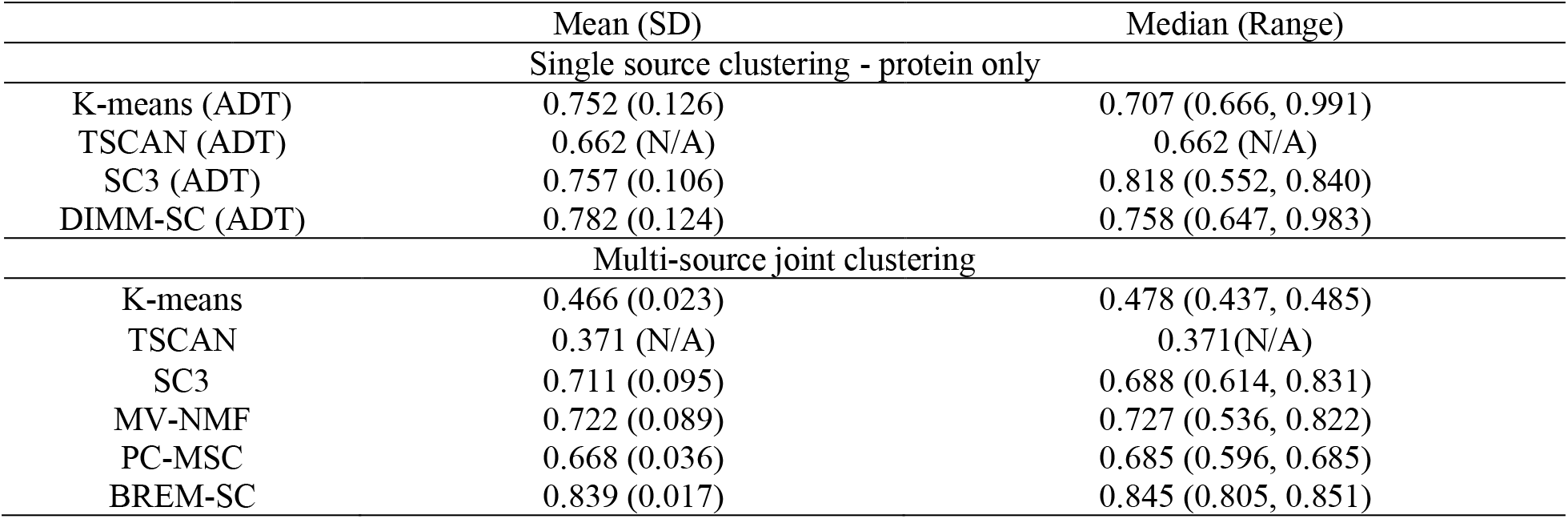
Performance (ARI) of clustering analysis for the subset of human PBMC real dataset with the removal of three protein markers (CD8A, CD16, CD127)

## DISCUSSION

When analyzing data from different data sources, ensemble clustering may be considered to integrate the separate clusters and determine an overall partition of cells that agrees the most with the source-specific cluster. However, most of ensemble clustering methods assume that the separate clusters are known in advance and do not inherently model the uncertainty (22). At the other extreme, a joint analysis that ignores the heterogeneity of the data may not capture important features that are specific to each data source. A fully integrative clustering approach is necessary to effectively combine the discriminatory power from transcriptome and protein measurements.

However, there are several noticeable limitations of this method. First, BREM-SC uses a computationally intensive MCMC algorithm, which is roughly linear with the number of cells and genes. In practice, it may result in a high computational cost when applying to large datasets (e.g., >10 K cells). To increase speed, BREM-SC has been implemented in R/Rcpp to accommodate data in relatively large scale. We further developed two alternative approaches to reduce running time. The first approach utilizes a naïve “block-wise Gibbs sampling” by which we first divide cells into multiple groups and use sequential Gibbs sampling within each group when updating cell-specific random effects. The benefit of this approach is to allow parallel computing with multiple CPUs, each for a block, to speed up this time-consuming step. Such an implementation may cause trouble under certain circumstances, for instance, when the covariates are highly correlated. However, under our model setting we expect the algorithm should still converge with this implementation, although the increasing number of blocks (or CPUs) may lead to some loss of efficiency. To assess the validity of this approach as well as the effect of the number of CPUs on the clustering result, we ran an additional simulation study and compared the ARI of clustering results between BERM-SC with parallel setting and regular setting. We simulated 100 datasets under our proposed model, and the results of ARI are summarized in **Supplementary Table S1**. In general, the results are consistent with our expectation, that larger number of CPUs set in parallel computing would result in lower efficiency of the convergence of algorithm. However, such a loss in efficiency is only subtle, and can be simply remedied by increasing the number of MCMC draws.

The second approach is to simplify our method by removing the random effects and assume independency between two data sources (illustrated in Equation 1). By assuming each of the two data follows separate multinomial distribution with Dirichlet prior, we derive the joint likelihood of two data and use E-M algorithm (23) to update parameters. We named this approach jointDIMM-SC, as it can be considered as a direct extension of DIMM-SC. Because of the fast speed of E-M algorithm, jointDIMM-SC runs only within several minutes for a dataset that BREM-SC would run for hours. This method is useful when there are in fact relatively low among-cell variabilities. We compared the performance of BREM-SC and jointDIMM-SC on the two real PBMC datasets. The estimated standard deviation of random effects for the public PBMC data is 0.75, and for the in-house data is 1.22. **Supplementary Table S2** summarizes the ARI of each method performed on each dataset, by which we observed that the performance of jointDIMM-SC is always worse than BREM-SC, and such a difference is larger when the estimated variance of random effects is larger. **Supplementary Figure S3** and **Figure S4** show the clustering results of jointDIMM-SC on public and in-house PBMC datasets, respectively, on UMAP plots. Although jointDIMM-SC performs worse than BREM-SC, still it could beat most of the competing methods currently available.

Both approaches have been implemented in our method as an optional choice. In addition to these, potential speed-up methods could include using graphics processing unit. Another limitation is that BREM-SC model ignores the measurement errors and uncertainties buried in count matrices. Multiple factors such as drop-out event, mapping percentage and PCR efficiency are not considered in the current model. These limitations can be largely overcome by extending the method. We will explore these directions in the near future.

In summary, we provide a novel statistical method, BREM-SC, for joint clustering scRNA-Seq and CITE-Seq data, which facilitates rigorous statistical inference of cell population heterogeneity. Our model-based joint clustering method can be readily extended to accommodate more than two data sources or multi-source data from other fields. We are confident that BREM-SC will be highly useful for the fast-growing community of large-scale single cell analysis.

## Supporting information

All Supplementary Data

## AVAILABILITY

The in-house CITE-seq data will be deposited into Gene Expression Omnibus (GEO). An R package of BREM-SC will be available on GitHub.

## ACCESSION NUMBERS

The published human PBMC CITE-Seq dataset that supports the finding of this study can be downloaded from 10X Genomics website (https://support.10xgenomics.com/single-cell-gene-expression/datasets/3.0.0/pbmc_10k_protein_v3). The in-house CITE-seq data will be deposited into Gene Expression Omnibus (GEO) and available on GitHub.

## SUPPLEMENTARY DATA

Supplementary Data are available at NAR online.

