## Supplementary material for "BREM-SC: A Bayesian Random Effects Mixture Model for Joint Clustering Single Cell Multi-omics Data": All Supplementary Data

**Supplementary Methods - Details of Gibbs Sampling**

We proposed a general Bayesian framework for estimation. We used Gibbs sample to iteratively update $z_j$, $\alpha_{i(k)}^{(1)}$, $\alpha_{d(k)}^{(2)}$ and $b_j$. Specifically, we used random walk Metropolis within Gibbs to iteratively update $b_j$, $\alpha_{i(k)}^{(1)}$ and $\alpha_{d(k)}^{(2)}$.

For a given cell $j$, the conditional distribution for $z_j$ follows a Multinomial distribution, where

$$P(z_j = k) = \frac{1}{constant}\left(\prod_{i=1}^{G}\frac{\Gamma\left(x_{ij}^{(1)} + \alpha_{i(k)}^{(1)}b_j\right)}{\Gamma\left(\alpha_{i(k)}^{(1)}b_j\right)}\right)\frac{\Gamma\left(\left|\boldsymbol{\alpha}_{(\boldsymbol{k})}^{(\mathbf{1})}b_j\right|\right)}{\Gamma\left(T_j^{(1)} + \left|\boldsymbol{\alpha}_{(\boldsymbol{k})}^{(\mathbf{1})}b_j\right|\right)}$$

$$\left(\prod_{d=1}^{D}\frac{\Gamma\left(x_{dj}^{(2)}+\alpha_{d(k)}^{(2)}b_j\right)}{\Gamma\left(\alpha_{d(k)}^{(2)}b_j\right)}\right)\frac{\Gamma(|\boldsymbol{\alpha}_{(\boldsymbol{k})}^{(\mathbf{2})}b_j|)}{\Gamma(T_j^{(2)}+|\boldsymbol{\alpha}_{(\boldsymbol{k})}^{(\mathbf{2})}b_j|)}\pi_k.$$

where the normalization constant is:

$$\sum_{k=1}^{K}\left\{\left(\prod_{i=1}^{G}\frac{\Gamma\left(x_{ij}^{(1)}+\alpha_{i(k)}^{(1)}b_j\right)}{\Gamma\left(\alpha_{i(k)}^{(1)}b_j\right)}\right)\frac{\Gamma(|\boldsymbol{\alpha}_{(\boldsymbol{k})}^{(\mathbf{1})}b_j|)}{\Gamma(T_j^{(1)}+|\boldsymbol{\alpha}_{(\boldsymbol{k})}^{(\mathbf{1})}b_j|)}\left(\prod_{d=1}^{D}\frac{\Gamma\left(x_{dj}^{(2)}+\alpha_{d(k)}^{(2)}b_j\right)}{\Gamma\left(\alpha_{d(k)}^{(2)}b_j\right)}\right)\frac{\Gamma(|\boldsymbol{\alpha}_{(\boldsymbol{k})}^{(\mathbf{2})}b_j|)}{\Gamma(T_j^{(2)}+|\boldsymbol{\alpha}_{(\boldsymbol{k})}^{(\mathbf{2})}b_j|)}\pi_k\right\}.$$

For a given gene $i$ and cell type $k$, the conditional log likelihood for $\alpha_{i(k)}^{(1)}$ is

$$logP\left(\alpha_{i(k)}^{(1)}\middle|\ldots\right) \propto \sum_{j=1}^{C} I(z_j = k)\log\left\{\left(\frac{\Gamma\left(x_{ij}^{(1)}+\alpha_{i(k)}^{(1)}b_j\right)}{\Gamma\left(\alpha_{i(k)}^{(1)}b_j\right)}\right)\frac{\Gamma(|\boldsymbol{\alpha}_{(\boldsymbol{k})}^{(\mathbf{1})}b_j|)}{\Gamma(T_j^{(1)}+|\boldsymbol{\alpha}_{(\boldsymbol{k})}^{(\mathbf{1})}b_j|)}\right\}.$$

Similarly, for a given ADT marker $d$ and cell type $k$, the conditional log likelihood for $\alpha_{d(k)}^{(2)}$ is

$$logP\left(\alpha_{d(k)}^{(2)}\middle|\ldots\right) \propto \sum_{j=1}^{C} I(z_j = k)\log\left\{\left(\frac{\Gamma\left(x_{dj}^{(2)}+\alpha_{d(k)}^{(2)}b_j\right)}{\Gamma\left(\alpha_{d(k)}^{(2)}b_j\right)}\right)\frac{\Gamma(|\boldsymbol{\alpha}_{(\boldsymbol{k})}^{(\mathbf{2})}b_j|)}{\Gamma(T_j^{(2)}+|\boldsymbol{\alpha}_{(\boldsymbol{k})}^{(\mathbf{2})}b_j|)}\right\}.$$

Finally, for a given cell $j$, we have the conditional log likelihood for $b_j$ as:

$$logP(b_j|\ldots) \propto \sum_{k=1}^{K} I(z_j = k))\ log$$

$$\left\{\left(\prod_{i=1}^{G}\frac{\Gamma\left(x_{ij}^{(1)}+\alpha_{i(k)}^{(1)}b_j\right)}{\Gamma\left(\alpha_{i(k)}^{(1)}b_j\right)}\right)\frac{\Gamma\left(\left|\boldsymbol{\alpha}_{(\boldsymbol{k})}^{(\mathbf{1})}b_j\right|\right)}{\Gamma\left(T_j^{(1)}+\left|\boldsymbol{\alpha}_{(\boldsymbol{k})}^{(\mathbf{1})}b_j\right|\right)}\left(\prod_{d=1}^{D}\frac{\Gamma\left(x_{dj}^{(2)}+\alpha_{d(k)}^{(2)}b_j\right)}{\Gamma\left(\alpha_{d(k)}^{(2)}b_j\right)}\right)\frac{\Gamma\left(\left|\boldsymbol{\alpha}_{(\boldsymbol{k})}^{(\mathbf{2})}b_j\right|\right)}{\Gamma\left(T_j^{(2)}+\left|\boldsymbol{\alpha}_{(\boldsymbol{k})}^{(\mathbf{2})}b_j\right|\right)}\right\} -logb_j - \frac{(logb_j)^2}{2\sigma_b^2}.$$

**Table S1.** Performance (ARI) of BREM-SC with parallel computing setting

| | Mean (SD) | Median (Range) |
|---|---|---|
| One CPU | | |
| 100 MCMC draws | 0.843 (0.108) | 0.898 (0.661, 0.974) |
| Ten CPUs | | |
| 100 MCMC draws | 0.830 (0.112) | 0.896 (0.631, 0.974) |
| 200 MCMC draws | 0.841 (0.111) | 0.903 (0.660, 0.973) |
| Twenty CPUs | | |
| 100 MCMC draws | 0.815 (0.112) | 0.884 (0.641, 0.961) |
| 200 MCMC draws | 0.828 (0.117) | 0.905 (0.655, 0.980) |

**Table S2.** Performance (ARI) of jointDIMMSC compared to BREMSC on two real CITE-seq datasets

| | Mean (SD) | Median (Range) |
|---|---|---|
| Public human PBMC ($\widehat{\sigma_b} = 0.75$) | | |
| BREM-SC | 0.737 (0.125) | 0.713 (0.556, 0.868) |
| jointDIMM-SC | 0.650 (0.107) | 0.658 (0.500, 0.851) |
| In-house human PBMC ($\widehat{\sigma_b} = 1.22$) | | |
| BREM-SC | 0.857 (0.048) | 0.874 (0.749, 0.895) |
| jointDIMM-SC | 0.767 (0.054) | 0.773 (0.638, 0.824) |

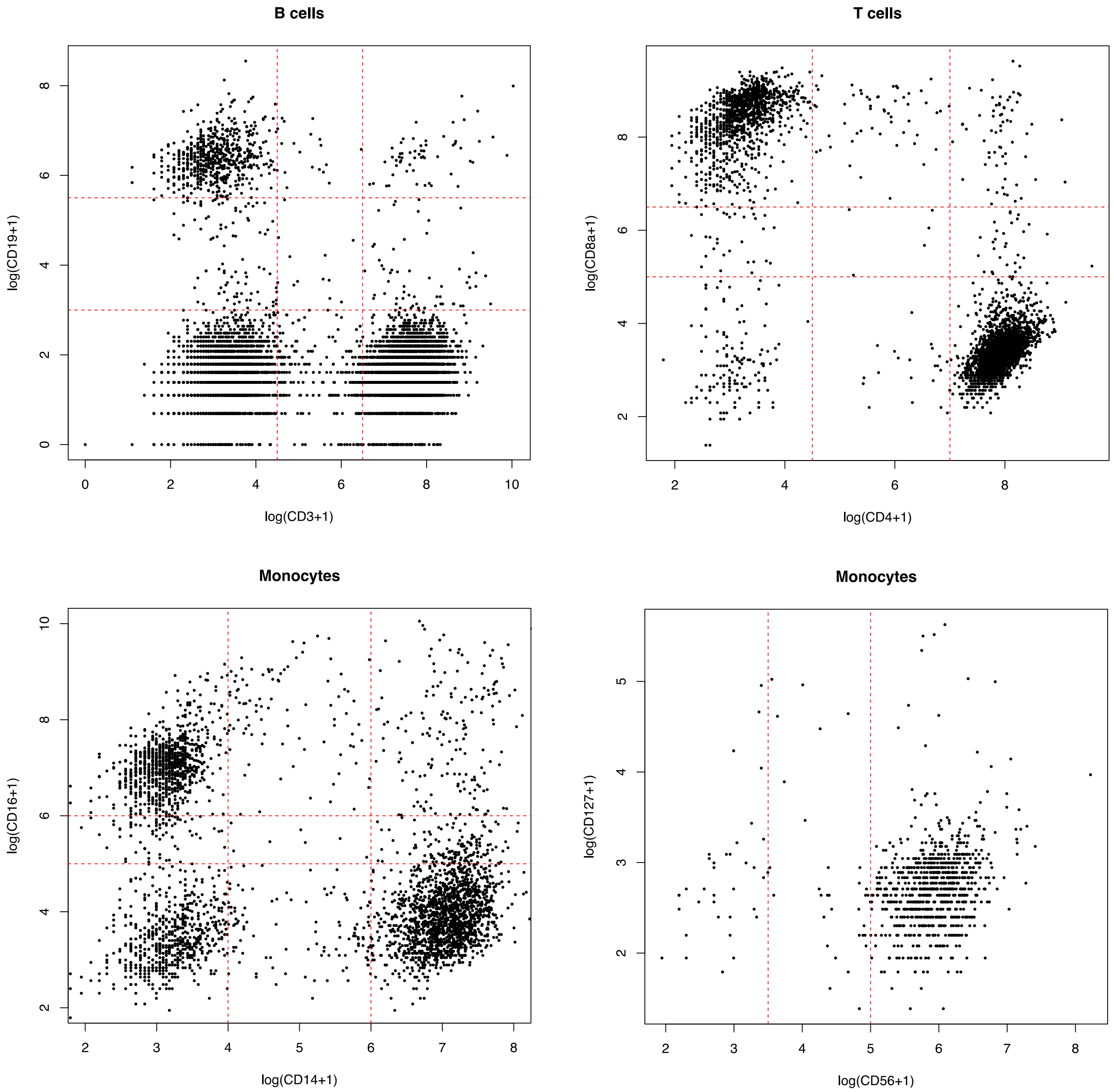


**Figure S1.** Scatter plot of cells illustrating how to get the approximated truth in 10X public human PBMC dataset.

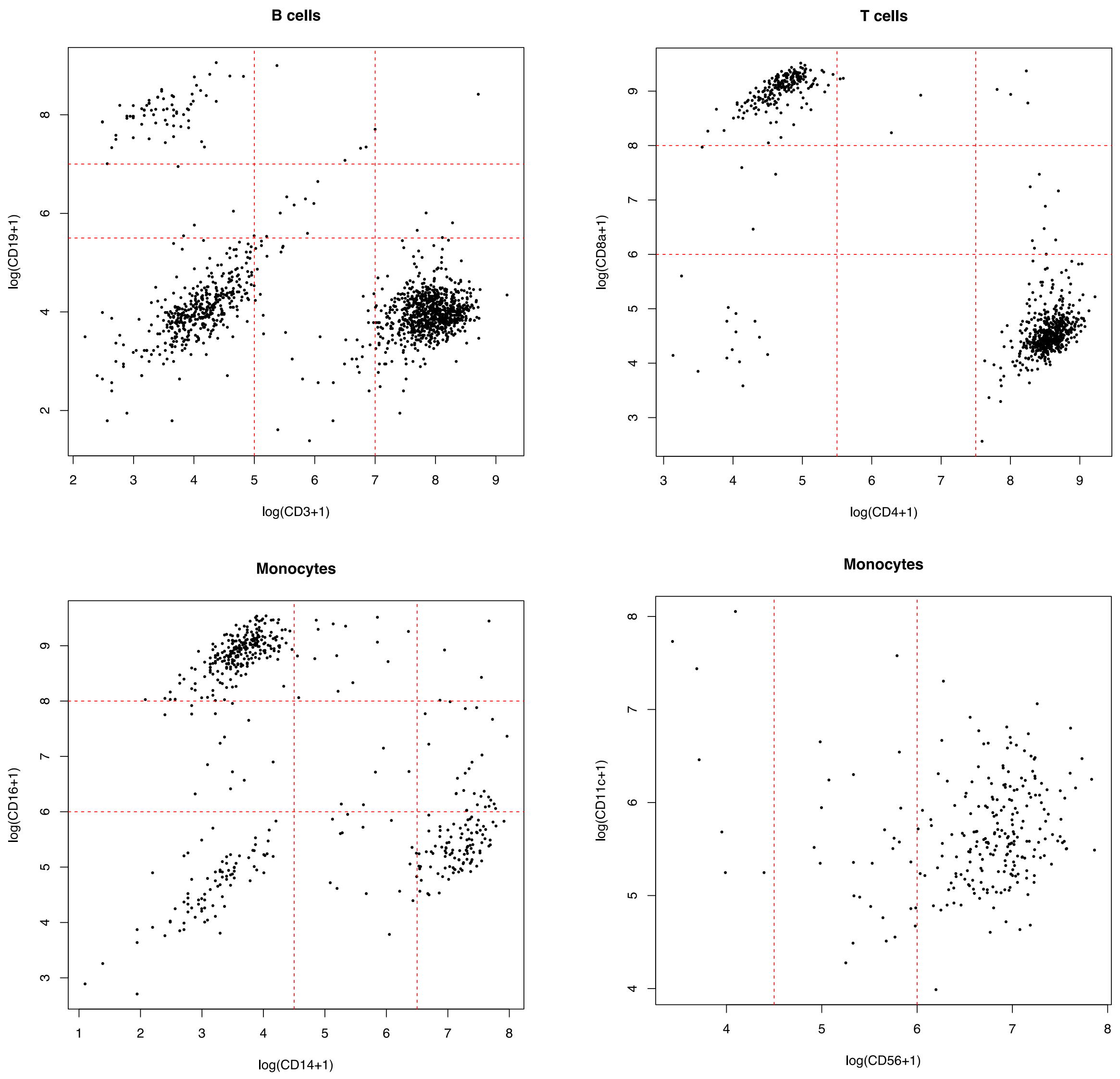


**Figure S2.** Scatter plot of cells illustrating how to get the approximated truth for in-house human PBMC dataset

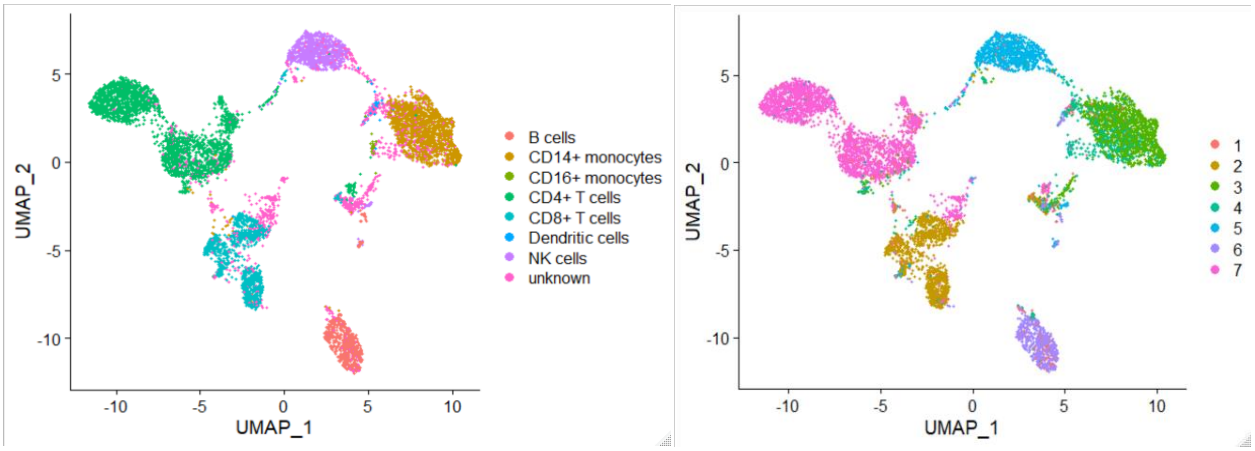


**Figure S3.** The performance of jointDIMM-SC for 10X public human PBMC CITE-Seq dataset. The UMAP projection of cells are colored by the ground truth (left) and jointDIMM-SC clustering results (right).

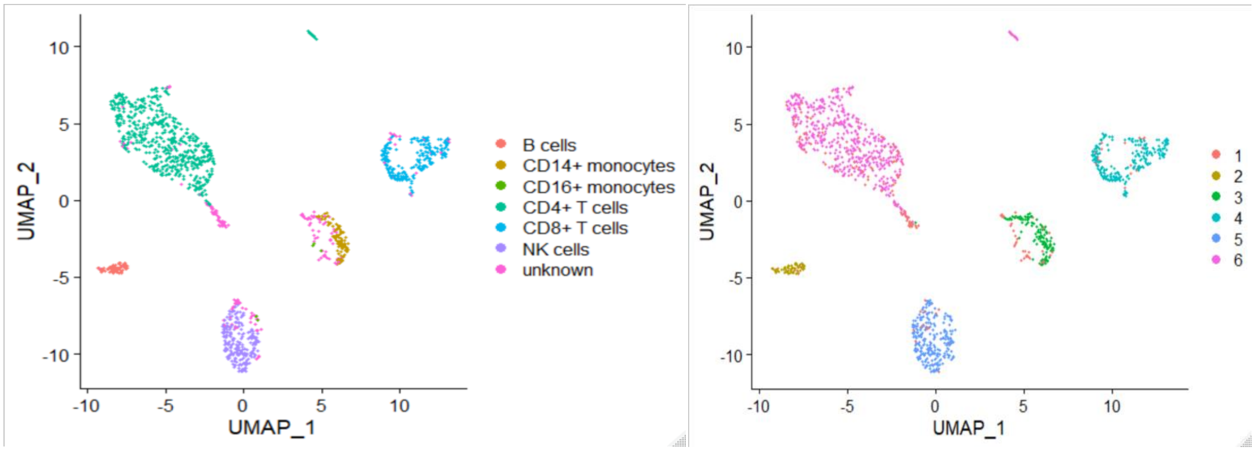


**Figure S4.** The performance of jointDIMM-SC for in-house human PBMC CITE-Seq dataset. The UMAP projection of cells are colored by the ground truth (left) and jointDIMM-SC clustering results (right).
